# Threatened species bycatch hotspots in the tropical Atlantic Ocean

**DOI:** 10.64898/2026.07.18.737235

**Authors:** Clara Lerebourg, Lucy Arnaud, Philippe S. Sabarros, David March, Mariana Travassos Tolotti, Christine N. Meynard, David M. Kaplan

## Abstract

Predicting bycatch patterns in tropical tuna purse seine fisheries remain challenging due to low variance explained by existing models, limiting their use for management purposes. To address this limitation, we combined ecological network analyses and community detection methods to characterize recurring bycatch assemblages in tropical tuna purse seine fisheries. Ecological network analyses consistently revealed two key bycatch communities: one dominated by bony fishes with low conservation concern, and another by IUCN-listed threatened species (NT-CR; sharks, rays, sea turtles). By focusing on sets with high numbers of threatened species, we identified bycatch hotspots near Mauritania, Gabon and Angola. These hotspots result from seasonal fishing patterns and specific bycatch compositions, with specific composition in terms of sex and age, compared to the rest of the Atlantic. We suggest that the Mauritanian region is a potential candidate for shark nursery and sea turtle foraging grounds. These insights can guide the development of bycatch reduction strategies based on predictable spatio-temporal or community-specific features and provide a basis for targeted management actions.

## Introduction

The incidental capture of non-target species, known as bycatch, has become a major and contentious issue in tropical tuna purse-seine (PS) fisheries worldwide due to the use of drifting fish aggregating devices (dFADs; [1]). Tropical tuna PS fisheries rely heavily on dFADs, which are human-made floating objects deployed by fishers that drift freely for long periods in the open ocean [2]. DFADs make use of the natural behavior of tropical tunas to aggregate around floating structures, thereby greatly facilitating catch of tuna schools [3]. Fishing sets on aggregated schools typically have higher bycatch levels compared to sets on schools not associated with floating objects (i.e., free-swimming tuna schools, FSC; [4]. While these devices facilitate the capture of the three main tropical tuna species: skipjack (*Katsuwonus pelamis*), yellowfin (*Thunnus albacares*) and bigeye tuna (*Thunnus obesus*), they also attract a range of other species, leading to increased catch of unwanted species called bycatch [4; 5]. In the tropical eastern Atlantic Ocean, the European PS fleet, predominantly composed of French and Spanish vessels, has been the main surface fishing fleet in the tropical region since the 1960s [6]. This fleet accounts for 70% of the PS tuna caught in the region, with French and Spanish vessels operating in overlapping regions of the eastern tropical Atlantic Ocean [7]. The main source of accurate PS bycatch data comes from onboard human observer records [8]. Observer coverage of fishing trips and effort has reached approximately 100% of the fishing effort of French vessels in the Atlantic since 2014. This dataset provides a wide range of information on bycatch species. Some bycatch species tend to be concentrated in specific areas that can be considered biodiversity hotspots, i.e. regions characterized by a high number of rare or threatened species [9], such areas represent priority targets for spatial or temporal mitigation measures aimed at reducing overall conservation risk. Some Atlantic coastal regions have already been identified as nursery or breeding grounds (e.g., for loggerhead turtles in the vicinity of Cape Verde or near Mauritania; [10; 11]), but a large-scale understanding of bycatch hotspots in the tropical Atlantic Ocean is lacking due to the paucity of bycatch data, highlighting the potential value of PS observer data to understand the dynamics of bycatch species.

The use of statistical models to estimate unobserved bycatch is an effective approach to estimating total bycatch rates [12]. However, at the scale of a single fishing set, predictability remains low, explaining only between 3% and 15% of the variance of four relatively common bycatch species (e.g., silky sharks; [12]). Even when data are aggregated spatially or temporally, large uncertainties remain, although significant trends may emerge. This poor predictive capacity limits the effectiveness of adaptive management measures (such as dynamic marine protected areas, [13]. To overcome these limitations, new approaches need to be explored. One such approach is a community analysis that may allow for a more comprehensive understanding of species assemblages, their interactions and how they collectively respond to pressures. Recent studies using modelling approaches have highlighted the importance of community structure in species distributions [14; 15]. This approach reveals key ecological patterns, and helps identify bycatch hotspots over space and time, providing a stronger foundation for conservation and sustainable fishing strategies.

The aim of this study was to analyze bycatch rates of French tropical tuna PS vessels to determine community structure and the spatial and temporal distribution of bycatch hotspots. First, we identified the most caught species by French purse seiners and examined the percentage of catches made with dFADs to highlight the importance of dFADs in bycatch. We then analyzed the community structure of bycatch species, exploring the relationships between the species. Finally, we analyzed the hotspots of threatened and IUCN-listed species, as defined by large bycatch events, and investigated potential biological and ecological reasons for their occurrence and distribution. These analyses provide a better understanding of the spatial and temporal dynamics of PS bycatch, as well as a methodological template applicable to pelagic fisheries worldwide.

## Results

To investigate whether community-based approaches can improve our understanding of bycatch patterns, we first quantified the relative importance of species and their association with dFADs. We then examined species co-occurrence patterns to identify recurrent bycatch assemblages. Finally, we focused on the assemblage dominated by threatened species to assess whether its occurrence was spatially structured and associated with persistent bycatch hotspots.

### 1. Dominant bycatch species and percentage caught on dFADs

Bycatch in the Atlantic PS fishery was overwhelmingly dominated by species associated with dFADs, while threatened species represented a much smaller but distinct component of the catch. It was composed primarily of five major taxonomic groups: billfishes, other bony fishes (i.e., excluding tunas and billfishes), rays, sharks and turtles. Among them, 12 species are found on the IUCN red list, with conservation statuses ranging from near threatened to critically endangered (**Table S1**). DFADs played a central role in PS capture of these species, representing most catches for 28 of 33 species and 9 of 12 threatened species (our dataset from French fleet has 8,635 sets on dFADs (54%) and 7,353 sets on FSC (46%); **Table S1, S2, Fig. S1**). Other bony fish represented the vast majority of bycatch, with 4,369,151 individuals recorded, 99% of them caught using dFADs, at an average rate of 505 individuals per set. In contrast, threatened species occurred at much lower abundances, although sharks remained relatively common, accounting for 71% of the 26,308 individuals and an average of 2.15 individuals caught per dFAD set. Of the 1,549 sea turtles recorded in observations, 75% were caught in dFAD sets, though at a lower rate of 0.14 individuals per set (note that almost all turtles are released alive; [5]). Rays and billfishes were also frequently associated with dFADs, which represent 47% and 37% of their total PS catches with average capture rates of 0.06 and 0.3 individuals per set, respectively (**Table S1, S2, S3 and S4**).

### 2. Community structure

The predominance of species associated with drifting FADs raised the question of whether these bycatch species formed recurring species assemblages rather than constituting a random collection of taxa. Despite the taxonomic diversity of bycatch species, cluster analyses consistently revealed the existence of two major assemblages (**Fig. 1a**). The first assemblage (cluster A) was dominated by species IUCN-listed threatened species (NT-CR). In contrast, the second assemblage (cluster B) was composed primarily of “dFAD-associated” species (i.e., species typically found in large quantities associated with dFADs; [16]). Species belonging to cluster B accounted for most observed bycatch events, with 81% of fishing sets (6,821 of 8,400 sets) consisting exclusively of individuals from cluster B. Species belonging exclusively to cluster A occurred alone in only 1.2% of sets (101 of 8,400 sets), primarily in coastal waters off Gabon and Angola (**Fig. S2**). The dendrogram and NMDS provided a good representation of species relationships (stress = 0.16; RMSE = 0.4%).

**Figure 1:**
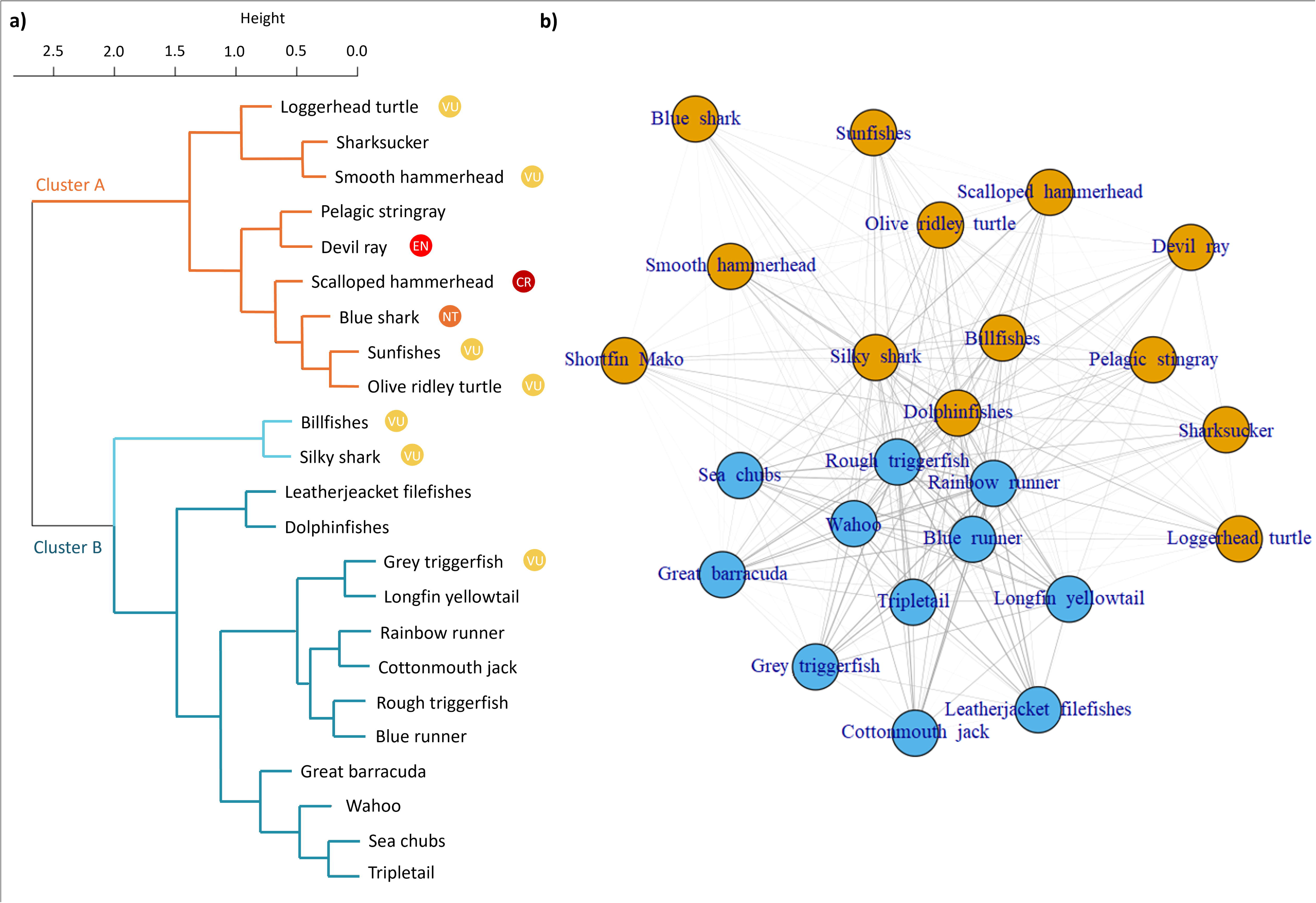
Community structure of bycatch species. Dendrogram of bycatch species and IUCN conservation status based on NMDS ordination (stress = 0.16) and Jaccard dissimilarity computed from a species-by-set presence–absence matrix (a), and co-occurrence network constructed from the same matrix (b; modularity = 0.09), to explore community structure and species associations. In panel (a), height represents the clustering distance (Jaccard dissimilarity) at which species or clusters are merged.

The network of co-occurrence analysis (**Fig. 1b**) obtained similar results to those of the dendrogram, with two groups composed of largely the same species as Clusters A and B identified. Notably, dolphinfishes, silky sharks and billfishes were included in the group of threatened species in the co-occurrence results. Although the Louvain algorithm identified two groups, the low modularity of the network (0.09) indicates weak community separation. This lower level of structure may partly arise because the network analysis was performed directly on the raw co-occurrence data, whereas the dendrogram was derived from the NMDS representation of species relationships, which reduces noise and emphasizes the dominant patterns in the data. Nevertheless, the species composition of the two groups was broadly consistent with the dendrogram and NMDS analyses. Together, these analyses indicate that Atlantic PS bycatch is structured around two recurrent assemblages: a widespread dFAD-associated community and a less common but distinct assemblage dominated by threatened species with dolphinfishes, silky shark and billfishes in some sense straddling the two groups.

### 3. Hotspot and distribution of bycatch species

The identification of a distinct community dominated by threatened species prompted us to investigate whether these species were also spatially clustered. We therefore focused on fishing sets characterized by high catches of threatened species to determine whether this assemblage exhibited spatial aggregation. Sets with high threatened species catches are clustered in four zones (**Fig. 2a**): the Cape Verde Basin off Mauritania (16° to 21°N - 21°W to 16°W), coastal waters off Gabon, between Cape Lopez and the islands of São Tomé (0° to 4°S - 10°E to 7°E), the Angolan Exclusive Economic Zone around the Cuanza River (13° to 7°S - 10°E to 14°E), and off the Guinea-Sierra Leone area (0° to 10°N - 21°W to 10°W). Although all four regions exhibited elevated catches of threatened species, only the Mauritanian, Gabonese and Angolan regions displayed species compositions that differed markedly from those observed across the broader Atlantic PS fishery (**Fig. 3a; Table S6**). By contrast, species composition in the Guinea–Sierra Leone region closely resembled that of the wider Atlantic despite locally high catch rates. According to Reid’s [9] definition of biodiversity hotspots, where a hotspot is characterized by a high concentration of rare or threatened species with a high risk of extinction, Mauritania, Gabon and Angola therefore emerged as the principal bycatch hotspots for threatened species in the tropical Atlantic Ocean.

**Figure 2:**
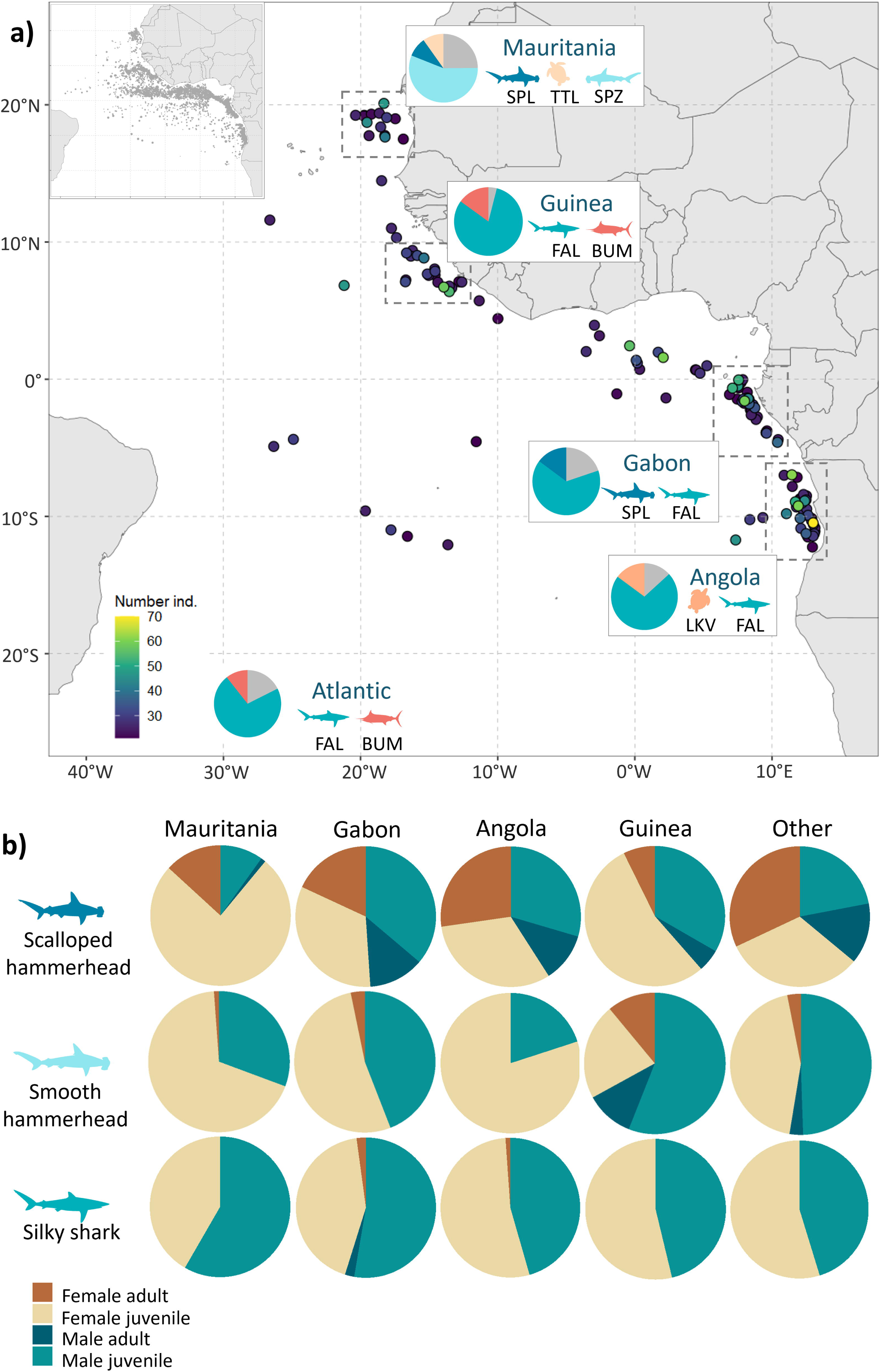
Hotspots of threatened-species bycatch. Bycatch hotspots of threatened species in the Eastern Atlantic with top species per hotspot (a; **Table S1**) and breakdown by sex and life-stage of the three top shark species caught in each hotspot and the Atlantic overall (b). In (a), only sets with >20 species individuals are shown and numbers of threatened species per set are limited to the range 20-70 to exclude a single outlier with 190 individuals (185 silky sharks, FAL, and 5 scalloped hammerhead, SPL) located near Gabon for a better visualization. Gray areas in pie charts in (a) represent other threatened species present in the area that individually represent less than 10% of the threatened species composition. Juvenile and adult life stages were discriminated based on the size-at-maturity (L50, **Table S5**). We then calculated the percentage of the different sex-maturity categories for each species. Only the three species shown had more than 30 individuals per hotspot. Detailed results of the species composition of the hotspots are available in **Table S6** (a) and percentages of adult and juvenile individuals, Chi2 results are available in **Table S7** (b), and contingency table is available in **Table S8**.

**Figure 3:**
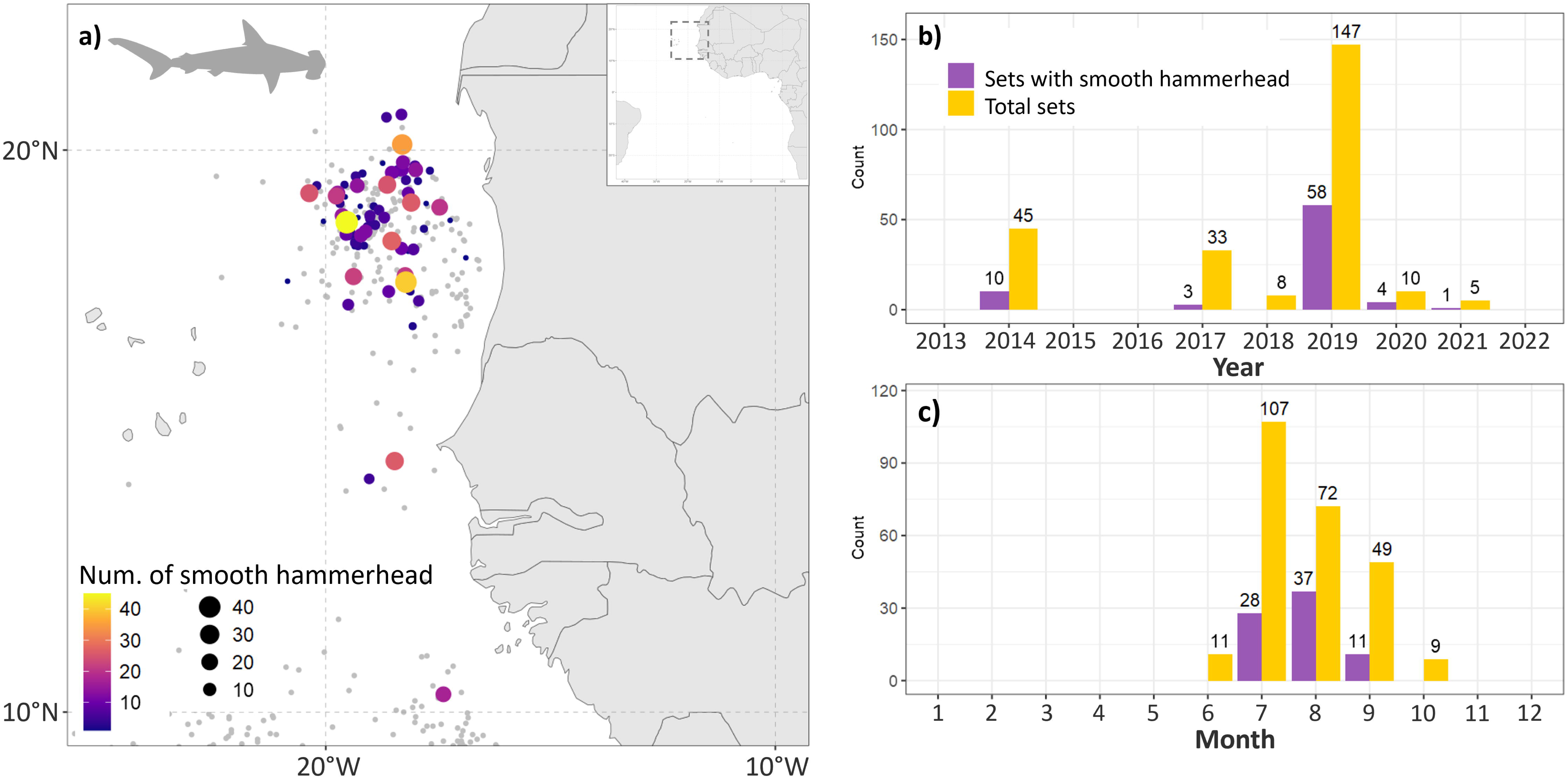
Smooth hammerhead distribution patterns. Spatial and temporal distribution of smooth hammerhead (SPZ). Map (a) illustrates the locations of smooth hammerhead captures in Mauritania. The colour and size of the points represent the relative importance of each capture set. The presence of grey points indicates sets that occurred in the region but did not result in the capture of any smooth hammerhead individuals. Graphs (b) and (c) illustrate the number of sets with SPZ by month and by year, respectively, along with the total number of sets conducted in Mauritania.

Each hotspot was characterized by a distinct assemblage of threatened species and demographic composition (**Fig. 2b; Tables S9–S11**). Across the Atlantic Ocean, silky sharks and billfishes were the most frequently observed threatened species in dFAD-associated catches, while silky sharks and smooth hammerheads were the most commonly captured shark species. Juveniles dominated catches of both species, representing 99% and 97% of individuals, respectively (**Table S7**). The Mauritanian hotspot was characterized by high catches of smooth hammerheads, scalloped hammerheads and loggerhead turtles (**Fig. 2a; Tables S7 and S9**). Juveniles accounted for more than 97% of shark catches, except for scalloped hammerheads, for which juveniles represented approximately 78% of individuals. Female juveniles were consistently overrepresented in both hammerhead species, whereas males occurred at lower frequencies. The Gabon hotspot exhibited a different species composition, dominated by silky sharks, scalloped hammerheads and smooth hammerheads, together with smaller numbers of billfishes, grey triggerfish and olive ridley turtles (**Fig. 2a; Tables S7 and S10**). Although juvenile sharks remained dominant, their proportion was generally lower than in Mauritania. Demographic composition also differed, with smooth hammerheads containing more juvenile males, scalloped hammerheads showing higher proportions of adult males, and silky sharks being predominantly male across both juvenile and adult stages. The Angolan hotspot was dominated by silky sharks, grey triggerfish and olive ridley turtles (**Fig. 2a; Tables S7 and S11**), with occasional catches of billfishes, scalloped hammerheads and devil rays. Nearly all smooth hammerheads were juveniles, and females accounted for 76% of individuals.

Statistical comparisons confirmed that demographic composition differed significantly among hotspots for all major threatened species examined (**Fig. 2b; Table S12**). For hammerhead sharks, differences in sex and maturity composition were significant but of moderate magnitude, with juvenile females being disproportionately represented in Mauritania relative to the other hotspots. Effect sizes were moderate for scalloped hammerheads and smaller for smooth hammerheads, indicating consistent but relatively limited demographic differentiation among regions. Silky sharks also exhibited significant spatial variation in demographic structure, with males predominating in Gabon across both juvenile and adult stages. However, effect sizes remained small, suggesting that the biological differences among hotspots were less pronounced than for hammerhead species despite strong statistical support. Additional analyses revealed significant spatial structuring for both rays and blue sharks (**Table S7 and S12**). Because several demographic categories contained low expected frequencies, Fisher’s exact tests were applied, with Monte Carlo simulations used for rays. Ray assemblages displayed moderate differences in demographic composition among hotspots, whereas blue sharks exhibited stronger demographic segregation. Confidence intervals could not be estimated for some sex and life stage composition because of sparse observations.

Long-term catch data reveal that the three most caught shark species, hammerheads and silky sharks, displayed year-round presence at regional hotspots, independent of month or year, when a minimum threshold of 30 sets were considered. In the Mauritania hotspots, smooth hammerheads were consistently recorded throughout the year (**Fig. 3**) and were more prevalent than elsewhere in the Atlantic. Scalloped hammerheads showed a similar year-round pattern in both Mauritania and Gabon but were more frequently encountered in the Gabon hotspot (**Fig. S3; Table S9 and S10**). Silky sharks also demonstrated continuous presence across years and months in Gabon suggesting an annual rather than seasonal presence (**Fig. S4**).

This temporal persistence occurred despite marked differences in regional fishing patterns (**Fig. S1**). Fishing effort was strongly concentrated between July and September in Mauritania (ρ = 0.88, indicating strong seasonality), occurred mainly between June and November in Gabon (ρ = 0.64), peaked between September and November in Angola (ρ = 0.89), and was more evenly distributed throughout the year in the Guinea–Sierra Leone region (ρ = 0.32, indicating weak seasonality). The recurrent occurrence of hotspot-associated species across these contrasting seasonal patterns suggests that the identified hotspots are associated with persistent ecological features rather than simply reflecting temporal concentrations of fishing effort.

## Discussion

The findings of this study highlight the heterogeneous spatial structure and community composition of dFAD bycatch, particularly in relation to vulnerable and endangered marine species. Of the 33 species identified as predominant in dFAD bycatch, 12 are listed as threatened by the IUCN, with conservation status ranging from near threatened to critically endangered. DFADs account for the majority of purse-seine captures among these species, though they very likely represent the minority of overall fisheries mortality. For example, hammerheads and turtles are likely caught at far higher rates in pelagic longlines and gillnets [17; 18]. The spatial distribution of threatened species highlighted four key bycatch hotspots in Mauritania, Guinea, Gabon, and Angola. In the case of Mauritania, the higher proportion of juvenile sharks relative to elsewhere in the Atlantic Ocean suggests that this hotspot may serve as a nursery or pupping ground for these species.

### 1. Catch on dFADs

In the Atlantic, dFADs are strategically deployed by tuna purse seiners along the West African coast, from Mauritania to Angola. The highest concentration of dFAD activity was recorded around the Gulf of Guinea and off the coasts of Gabon and Angola [19], which likely reflects the greater concentration of dFAD-associated fishing effort in these regions, resulting in a higher number of threatened species being recorded in the bycatch. Similar results on the distribution of shark species were found [6; 20]. Among a total of 4,405,146 individuals, 99.5% were caught using dFADs, in line with the literature [2; 5].

### 2. Clustering and its consequences

Our dendrogram analysis revealed two principal groups of PS bycatch species around dFADs. Though these results are consistent across two approaches to clustering species, the relatively weak separation of the network analysis, the large number of rare species and the prevalence of more common species throughout the fishing zone are indicative of PS bycatch species compositions being highly stochastic, with some consistent co-occurrence patterns, but no strong segregation into distinct ecological communities. Results are generally consistent with what is known about co-occurrence of epipelagic species in regions other than the tropical Atlantic Ocean. In the Pacific Ocean, silky shark and blue marlin not only share part of their habitat but also have dietary similarities [21]. Their piscivorous diets are dominated by tuna and both species are mainly observed near the surface. This brings them in line with dolphinfish, with whom they share the status of epipelagic species, restricted in their movements by the thermocline, with a similar diet of epipelagic prey [22; 23]. They also support these observations in the South Atlantic, with dendrograms showing a close grouping between silky sharks and blue marlins, with dolphinfish forming a slightly more distant, but still close subgroup [22]. Our results also highlight the clustering of threatened species, such as rays, sharks, turtles, and remoras, which can be explained by factors like distribution and diet. For instance, the devil ray and the pelagic stingray both feed on zooplankton and small fish [24; 25]. Additionally, the sicklefin devil ray has been observed hunting cooperatively with the common remora in the Azores archipelago [26]. Similarly, sharks, including scalloped and smooth hammerheads, as well as shortfin makos, primarily consume fish, crustaceans, and cephalopods, with squid being a key prey item [17; 21]. From a mechanistic perspective, the observed co-occurrence patterns may be attributed to the attraction of these species to dFAD-associated prey fields or to aggregation processes occurring at specific oceanographic features, such as fronts, where enhanced productivity leads to a concentration of forage species. dFADs themselves have been identified as drifting ecological hotspots, with the potential to increase local prey availability and facilitate multi-species aggregations across trophic levels.

### 3. Hotspots, biology and shark nursery

The four identified bycatch hotspots correspond to areas of high bycatch occurrence based on effort-standardized purse seine data. The first hotspot, off the coast of Mauritania (16° to 21°N, 21°W to 16°W), lies near the continental shelf edge, south-west of Banc d’Arguin, in a region rich in coastal pelagic fish. The literature shows that this area experiences consistently high concentrations of inorganic nutrients, chlorophyll-a, and primary production throughout the year, with peaks in winter [27], which are likely to support elevated trophic activity. The second hotspot, near the region of Guinea (0 to 10°N, 21°W to 10°W), is characterized by the Guinea dome, a distinctive thermocline structure that significantly enhances primary productivity. This is believed to support the development of the trophic chain [28; 29]. The third hotspot near Gabon (0° to 4°S, 10°E to 7°E), is influenced by a thermohaline front from June to September south of São Tomé [30], which supports a high biomass of chlorophyll-a, zooplankton, and micronekton, attracting species like tuna and marine species that can contribute to bycatch [28; 31; 32]. The fourth hotspot, off Angola (13° to 7°S, 10°E to 14°E), also aligns with areas of high productivity, with cold and nutrient-rich upwelled waters of the Benguela current to the south driving rapid phytoplankton growth which in turn sustains higher trophic levels [33]. These areas of high productivity often lead to the development of larger and more active fishing grounds, as the abundance of prey species attracts both target and non-target species populations. A complete accounting of the relationship between target and non-target species would, however, require data from multiple fishing gears, particularly gears such as longline and gillnets that are often the primary sources of fishing mortality for vulnerable species.

According to Heupel et al., [34], a marine area can be considered a shark nursery if it meets three criteria: sharks must be observed there more frequently than in surrounding areas (preference), they must stay there or return for long periods (residence), and the area must be used repeatedly over the years (consistency). While our results provide partial support for the preference criterion in Mauritania, particularly for smooth hammerhead sharks with a high proportion of females and juveniles, the absence of movement or residency data prevents full validation of nursery function. Therefore, this area should be considered a candidate nursery or pupping ground. The same criteria were applied to the scalloped hammerhead and the silky shark, but these species didn’t meet the preference criterion (as their main hotspot were respectively in Gabon and Angola). In a regional study based on artisanal fishery catch, Doherty et al., [35] found a biodiversity hotspot for sharks and rays in the waters of Congo immediately adjacent to the Gabon hotspot. They found a seasonal pattern in the distribution of species, with high abundance of scalloped hammerheads from August to December (and lower from February to July), of blue sharks in the middle of the year (May to September), while for silky shark’s catches remained constant throughout the year. These results are consistent with a seasonal structuring of shark populations in the region, although direct comparisons are limited by differences in fishing gear and sampling design. In addition, for silky sharks and scalloped hammerheads, Doherty et al., [35] caught mainly immature individuals and hypothesized the presence of nursery or pupping grounds in the EEZ of Congo. In Gabon, the scalloped hammerhead was the only species to meet the preference criterion, indicating frequent use of this area. However, because this species was not observed every month, it does not satisfy the consistency criterion. Finally, in Angola, silky shark is the most common threatened bycatch species, with a majority of young individuals caught, though the fraction of juveniles is not significantly different from elsewhere in the Atlantic Ocean, but this species was recorded every month and every year, fulfilling the consistency criterion. Nonetheless, assessment of the residence criterion requires additional data, such as tracking or recapture studies, which are not available for this study.

Similar patterns were observed for sea turtles. In the eastern Atlantic, major loggerhead turtle nesting colonies are located in Cape Verde, whereas important olive ridley nesting populations occur along the Gulf of Guinea, notably in Angola, Gabon and Ghana [36; 37]. Telemetry studies have shown that sea turtles can undertake large-scale movements between nesting and foraging areas, with their distribution partly influenced by ocean productivity patterns and environmental conditions [38]. Consequently, bycatch hotspots identified in Mauritania and Angola may overlap with important foraging habitats used by individuals originating from these regional nesting populations. This highlights that PS interactions occurring in these hotspots may have consequences extending beyond the local scale by affecting populations connected to distant breeding colonies. However, additional tracking and population-assignment studies would be required to quantify these links directly.

A comprehensive understanding of the relationship between target and non-target species, as well as the ecological significance of identified hotspots, would require integrating data from multiple fishing gears. In particular longlines and gillnets, which are known to account for a substantial proportion of mortality in vulnerable species such as sharks, rays and turtles, may exhibit different spatial patterns. Consequently, the hotspots identified here should be interpreted as PS interaction hotspots, rather than absolute indicators of species distribution or total fishing risk.

### 4. Management implications and research priorities

The identification of spatially structured bycatch hotspots allows for the development of targeted and operational management measures. Several concrete interventions can be derived from our results. The clear aggregation of threatened species in Mauritania, Gabon and Angola suggests that dynamic or seasonal spatial closures could be effective in reducing interactions between PS fisheries and vulnerable species, particularly when implemented during periods of peak occurrence. Other complementary mitigation measures, such as real-time hotspot avoidance programs, improved bycatch release protocols, or adaptive management based on near real-time species distribution forecasts, may also contribute to reducing bycatch while limiting disruptions to fishing activities.

For instance, in Mauritania where a high proportion of juvenile hammerhead sharks is observed, a seasonal closure during peak juvenile occurrence could reduce mortality in early life stages. In Gabon and Angola, where threatened shark species are consistently present, recurrent seasonal closures could be implemented. The key missing factor to accurately evaluate the effectiveness of such measures is a more complete understanding of bycatch and mortality across multiple fishing gears, particularly longlines and gillnets, which may account for a substantial proportion of mortality in threatened species. In addition, future work should assess the socioeconomic consequences of potential spatial or seasonal closures on fishing operations in order to balance conservation benefits with fishery viability.

Finally, Mauritania and potentially Gabon emerge as candidates for shark nursery zones. These areas could be prioritized for enhanced protection status though further validation via, for example, tagging and tracking studies should be a research priority before formal designation.

### 5. Data limitations and potential detection bias

A key limitation of this study is the uneven spatial distribution of PS effort data across the study region. Areas such as the EEZs of Togo, Benin, Nigeria, Cameroon, Equatorial and Congo lacked data due to an absence of fishing agreements for French PS, which constrains our ability to detect the presence of bycatch species in these regions. As a result, the absence of identified hotspots in these areas should not be interpreted as evidence of low bycatch risk, but rather as a potential artefact of the distribution of PS fishing effort (i.e. false negatives). Consequently, the hotspots identified here should be interpreted as areas of high observed interaction given available data, rather than exhaustive representations of bycatch risk across the tropical Atlantic. Future work should prioritize improving data coverage in underrepresented regions through the integration of data from multiple purse-seine fleets operating in the Atlantic Ocean, as well as from other fishing gears to better quantify spatial effort distribution and refine hotspot detection.

## Conclusion

Our study revealed that the twelve bycatch species classified as threatened by the IUCN that are regularly caught by French PS vessels in the tropical Atlantic Ocean stand out in terms of co-occurrence patterns and spatial distributions relative to other bycatch species. In particular, sets with important catches of these species are not homogeneously distributed throughout the fishing zone, but rather are concentrated in four hotspots, three of which are clearly distinct in bycatch species composition of threatened species relative to each other and the wider Atlantic Ocean: Mauritania, Gabon and Angola. The fourth hotspot, Guinea, has a similar species composition to the rest of the Atlantic fishing zone, with a distinct size-sex composition for endangered sharks. Several of these zones are candidate nursery areas, including Mauritania for scalloped hammerhead sharks and Gabon silky sharks. Collectively, these results provide considerable baseline information for structuring spatial management and bycatch mitigation efforts in the area. The key question for future research is the extent to which these zones identified based on PS bycatch observations can be used to define cross-fisheries management strategies to reduce bycatch of endangered pelagic megafauna. Our findings support the implementation of spatio-temporal management measures such as dynamic closures. These measures, combined with cross-fishery management approaches, offer a practical pathway to reduce bycatch of threatened pelagic species.

## Materials and Methods

### 1. Data collection

Data for this study were obtained from human observer programs on board French tuna purse seiners operating in the tropical Atlantic Ocean between 2013 and 2022. These programs include data collected through the European Union Data Collection Framework (DCF, EU Regulation 199/2008), the International Commission for the Conservation of Atlantic Tunas (ICCAT) Moratoria program, and the Observateur Commun Unique et Permanent (OCUP) program managed by the French producer organization ORTHONGEL [39]. All three data collection initiatives follow a common and standardized protocol [40]. Trained observers on board the vessels record essential information for each fishing set, including date, time, geographic coordinates, and estimate total bycatch per species in terms of number of individuals. Observers were trained using standardized identification guides, and sensitive species (e.g. sea turtles, sharks, rays, billfishes) were systematically photographed to allow post hoc validation. Individuals were measured following species-specific length protocols (e.g. total length for sharks, disc width for rays and curved carapace length for sea turtles) and sex is recorded when it can be determined (notably for sharks, rays and turtles), otherwise noted as indeterminate.

### 2. Selected species

To identify the species most commonly caught by purse seiners, we performed a selection process (**Fig. S5**). First, we analysed the total catch data to select the 40 species most frequently caught by French purse seiners between 2013 and 2022. For that we focused on catches made with dFADs, retaining only those species for which more than 100 individuals were caught, and excluding the catches made on tuna free-swimming schools (FSC; agregatings of tuna that are not associated with floating objects like dFADs). The threshold of 100 individuals was chosen to limit the influence of extremely rare observations that may introduce noise and unstable associations. Similarly, restricting the analysis to dFAD-associated sets allows focusing on a consistent fishing mode known to generate distinct bycatch assemblages, while avoiding confounding effects linked to different fishing strategies such as FSC sets. The dataset was then filtered, as explained in the following sections, to conduct clustering and co-occurrence network theory analyses, identification of hotspots and additional analyses. We acknowledge that this filtering excludes rare species, including some threatened taxa. To mitigate this potential bias, we conducted complementary analyses using a dedicated biological dataset focusing exclusively on threatened species, independently of their frequency of occurrence. Analyses were based on presence-absence data rather than abundance to reduce the influence of highly abundant species and better account for infrequent occurrences.

### 3. Community structure

To understand the structure and composition of bycatch communities as a function of species present and catch location, we conducted a detailed analysis of bycatch communities in fisheries. We started by constructing a presence-absence matrix, where each species was coded as present (1) or absent (0) in each set. This approach provided more reliable and robust results than standardizing species abundances by their maximum values, as it treated all species equally and avoided distortions caused by extreme values or highly abundant species. We only included species present in at least 1% of all the fishing operations, i.e. at least 84 occurrences out of 8,400 operations, to reduce noise associated with extremely rare species while retaining ecologically relevant taxa. Although the standard threshold for this type of selection is generally set at 5%, applying this value would have resulted in the exclusion of all threatened species from the dataset, which was not desirable given the conservation focus of the study. To assess the robustness of this threshold choice, we conducted a sensitivity analysis using alternative occurrence thresholds ranging from 0.5% to 5% of fishing operations. Lower threshold (0.5%) resulted in the inclusion of all species but increased potential noise associated with extremely rare taxa. Conversely, higher thresholds progressively excluded ecologically important and conservation-relevant species. The 1% threshold excluded 3 species (whitspotted filefish, blue shark et shortfin mako), the 2% threshold excluded 8 species (adding sharksucker, sunfishes, cottonmouth jack, devil ray, and loggerhead turtle) and the 5% species excluded 11 species (adding pelagic stringray, smooth and scalloped hammerhead). The 1% threshold provided a balanced compromise, minimizing noise (only 0.48% of individuals excluded) while retaining rare and conservation-relevant taxa. We also excluded operations where only one single species was caught, as these cases were considered outliers.

Once these data were cleaned and filtered, a Jaccard dissimilarity matrix was computed to measure differences in specific composition between sites using the ‘vegdist’ function in the ‘vegan’ R package [41]. This matrix was then used as the basis for multidimensional non-metric ordination (NMDS) using the Jaccard distance method, using the ‘metaMDS’ function in the ‘vegan’ R package. The NMDS analysis, configured to maximize fit and reduce stress (a measure of goodness of fit: stress <0.1 = good fit, <0.2 = acceptable fit, >0.3 = poor fit), projected the data into a three-dimensional space to visualize the relationships between sites in terms of compositional similarity. NMDS analyses were conducted by testing different dimensionalities (k = 2, 3 and 4). For each value of k, the algorithm was run with a maximum of 20 random starts (trymax = 20) to ensure convergence while maintaining reasonable computation of time. The selection of the optimal dimensionality was based on multiple criteria, including stress values, shepard plots, and goodness-of-fit diagnostics (**Fig. S6**). Although k = 4 yielded slightly lower stress values, the improvement compared to k = 3 was negligible, while k = 2 showed substantially higher stress. Therefore, k = 3 was retained as a compromise between model fit and interpretability. We determined the optimal number of clusters based on the presence–absence data. Two classification approaches were compared: an ascending hierarchical clustering method based on Ward’s algorithm, known for its robustness, and a k-means clustering method. Both approaches converged to the same result, indicating an optimal number of two clusters. Distances between sites were calculated and a dendrogram was constructed to visualize the clusters. To better understand these clusters, we examined the geographical distribution of the species within them.

The dataset used for NMDS analysis and dendrogram construction (where rows represent sites and columns represent species) was transposed into a co-occurrence matrix. Each element of this new matrix indicates the number of sites where two species co-occur. The diagonal values were set to zero to exclude co-occurrences of a species with itself.

To limit the bias associated with the frequency of occurrence of species, the co-occurrence matrix was normalized using the Szymkiewicz–Simpson overlap coefficient. This normalization was performed by dividing each element by the minimum of the total occurrences of the two corresponding species. Unlike the Jaccard index, which is strongly influenced by differences in species prevalence, the overlap coefficient is less sensitive to rare species and is therefore particularly suitable for ecological datasets containing many infrequent taxa. The normalized matrix was then used to construct a species network using the ‘graph_from_adjacency_matrix’ function in the ‘igraph’ R package [42]. This function transformed the co-occurrence matrix into an undirected weighted network. In this network, each node represents a species and each link, weighted by the strength of co-occurrence, symbolizes a relationship between two species (self-links were excluded to improve the readability of the network). The network was then used to identify species communities using the Louvain algorithm [43], which identifies groups of species based on the density of their connections within the network. Each community corresponds to a group of species that co-occur with each other more frequently than with other species. Finally, a graphical representation of the network was generated. The nodes were colored according to their community affiliation, while the thickness of the links reflected the intensity of co-occurrences between species. We then calculated the Louvain modularity, which is a key measure used to assess the quality of a community partition in a network. A modularity scores close to 1 indicates well-defined communities, while a score close to 0 indicates that the edges are randomly distributed.

### 4. Identification of hotspots and species biology

To identify bycatch hotspots, we analyzed the geographical distribution of threatened species and searched for areas where these species were disproportionately concentrated, following Reid’s [9] definition of biodiversity hotspots. To operationalize this concept, we first identified fishing sets associated with exceptionally high numbers of threatened individuals. A fishing set was classified as a hotspot event when it recorded ≥20 individuals belonging to one or more threatened species, corresponding to the 95th percentile of the observed distribution. Spatial aggregation of these hotspot events was then used to identify broader hotspot areas. We compared the resulting hotspot distributions using the 90th, 95th and 97th percentiles. The 95th percentile was retained as a compromise between capturing the major spatial patterns and focusing on the most extreme bycatch events. To account for spatial heterogeneity in sampling effort, hotspot identification was performed at the level of individual fishing sets (i.e. standardized sampling units). It is important to note that observer data were unavailable for certain regions (Togo, Benin, Nigeria, Cameroon, Equatorial Guinea and Congo). Consequently, the absence of identified hotspots in these areas should not be interpreted as evidence of low bycatch risk. In addition, other PS fleets may operate in some of these EEZs, meaning that bycatch interactions occurring there are not represented in the present dataset. Conversely, sampling effort was greater in Angola and Gabon, which may increase the probability of detecting hotspot events in these regions (**Fig. S7**).

We then looked in detail at the species present in these areas to distinguish the relative concentrations of males, females and juveniles. Of the 30,991 captured individuals of vulnerable species studied, 17,545 (56.6%) had biological data. We did not have the necessary biological information for sunfish (*Mola mola*), blue marlin (*Makaira nigricans*) and the grey triggerfish (*Balistes capriscus*) since sex cannot be externally determined by observers for these three species. Except for these three, we discriminated juvenile and adult life stages based on the size-at-maturity (L50) for each species available based on information available in the literature (**Table S5**). We then calculated the percentage of the different categories for each species: juvenile male, adult male, juvenile female and adult female. To assess whether there is a significant difference between the biological data in the different hotspots and the rest of the Atlantic, we performed Chi² tests of independence. This test is used to determine whether there is an association between two qualitative variables within a contingency table. To be valid, it must meet several conditions: (i) the data must be in count form, and the sample size must be large enough to limit statistical errors, with a headcount greater than 30, (ii) numbers must be greater than 5 in at least 80% of the cells in the table, and no theoretical number must be less than 1, and (iii) each observation must be independent and can belong to only one category. In addition to p-values, effect sizes were calculated using Cramer’s V to quantify the strength of the association between variables. For these analyses, we included counts for all non-hotspot zones in the Atlantic to compare hotspot compositions with those of the wider Atlantic.

Finally, seasonal patterns in PS fishing effort were quantified using circular statistics to account for the cyclic nature of monthly data. The fishing months were converted into angular values ranging from 0 to 2π. The mean direction (μ) was used to estimate when fishing activity peaks, while the length of the mean resultant vector (ρ) was used to quantify the intensity of seasonality, with values close to 1 indicating a strong temporal concentration and those close to 0 indicating a uniform distribution throughout the year. Circular statistics were calculated using the ‘circular’ package in R. Analyses were conducted separately for each hotspot.

## Acknowledgements

We thank Ob7, particularly Pascal Cauquil and the team handling bycatch data, for their assistance with data processing. We are also grateful to Fabien Forget, participants of the Common Oceans Tuna Bycatch workshop (Rome, 2025) and members of the REDUCE, MarineBeacon, and CiBRINA projects for their valuable feedback on our results. We also thank Julien Patras for his thorough review of the manuscript and his insightful comments.

## Author contributions

CL conducted all analyses and wrote the manuscript. The community structure analysis was carried out using R scripts and the internship report of LA, with support from CNM and DMK. DMK supervised the project alongside DMM, providing guidance throughout. All co-authors reviewed, edited and provided feedback on the manuscript.

## Data and Code availability

Due to the confidential nature in this study, access requests should be directed to the Ob 7 pelagic ecosystem observatory via their official website (https://www.ob7.ird.fr/) or by the email address.

## Github link

https://github.com/ClaraLerebourg/Lerebourg_et_al-Threatened-species-bycatch-hotspots-in-the-tropical-Atlantic-Ocean

