## Supplementary Information for "Threatened species bycatch hotspots in the tropical Atlantic Ocean"

#### Supplementary Tables

Table S1 : Table of the total observed number of bycatch individuals caught between 2013 and 2022 on drifting fish aggregating devices (dFADs), free-swimming tuna schools (FSC), and both (ALL), rate of bycatch individuals caught per dFAD set (no./ dFADset), and free-school set (no./ FSC set), and percentage caught of dFAD (% caught on dFAD). Observations represent 8635 sets on dFADs and 7353 on FSC.

| 2013-2022 | | | | | Number | | | | | |
|---|---|---|---|---|---|---|---|---|---|---|
| **Species group** | **Common name** | **FAO** | **Scientific name** | | **ALL** | **dFAD** | **FSC** | **no./dFAD set** | **no./ FSC set** | **% caught on dFAD** |
| Billfishes | Billfishes | BUM | *Makaira nigricans* | VU | 2,670 | 2,383 | 287 | 0.28 | 0.04 | 89.25 |
| | | SAI | *Istiophorus albicans* | / | 4,350 | 248 | 4102 | 0.03 | 0.56 | 5.70 |
| Other bony fishes | Leatherjacket filefish | ALN | *Aluterus scriptus* | LC | 390 | 390 | 0 | 0.05 | 0.00 | 100.00 |
| | | ALM | *Aluterus monoceros* | LC | 2,248 | 2,247 | 1 | 0.26 | 0.00 | 99.96 |
| | Rough triggerfish | CNT | *Canthidermis maculata* | LC | 1,230,707 | 1,229,973 | 734 | 142.44 | 0.10 | 99.94 |
| | Dolphinfishes | CFW | *Coryphaena equiselis* | LC | 120 | 116 | 4 | 0.01 | 0.00 | 96.67 |
| | | DOL | *Coryphaena hippurus* | LC | 55,106 | 54,741 | 365 | 6.34 | 0.05 | 99.34 |
| | Sharksucker | EHN | *Echeneis naucrates* | LC | 123 | 71 | 52 | 0.01 | 0.01 | 57.72 |
| | | REO | *Remora remora* | LC | 244 | 217 | 27 | 0.03 | 0.00 | 88.93 |
| | Great barracuda | GBA | *Sphyraena barracuda* | LC | 16,595 | 16,567 | 28 | 1.92 | 0.00 | 99.83 |
| | Whitespotted filefish | JKY | *Cantherhines macrocerus* | LC | 139 | 139 | 0 | 0.02 | 0.00 | 100.00 |
| | Sea chubs | KYP | *Kyphosus spp* | LC | 252 | 252 | 0 | 0.03 | 0.00 | 100.00 |
| | | KYS | *Kyphosus sectatrix* | LC | 13,072 | 13,015 | 57 | 1.51 | 0.01 | 99.56 |
| | Tripletail | LOB | *Lobotes surinamensis* | LC | 18,679 | 18,602 | 77 | 2.15 | 0.01 | 99.59 |
| | Sunfishes | MOX | *Mola mola* | VU | 324 | 122 | 202 | 0.01 | 0.03 | 37.65 |
| | | MRW | *Masturus lanceolatus* | LC | 146 | 80 | 66 | 0.01 | 0.01 | 54.79 |
| | | RZV | *Ranzania laevis* | LC | 128 | 107 | 21 | 0.01 | 0.00 | 83.59 |
| | Rainbow runner | RRU | *Elagatis bipinnulata* | LC | 792,554 | 790,986 | 1,568 | 91.60 | 0.21 | 99.80 |
| | Blue runner | RUB | *Caranx crysos* | LC | 2,079,235 | 2,078,124 | 1,111 | 240.66 | 0.15 | 99.95 |
| | Grey triggerfish | TRG | *Balistes capriscus* | VU | 8,688 | 8,660 | 28 | 1.00 | 0.00 | 99.68 |
| | Cottonmouth jack | USE | *Uraspis secunda* | LC | 13,112 | 13,112 | 0 | 1.52 | 0.00 | 100.00 |
| | Wahoo | WAH | *Acanthocybium solandri* | LC | 55,181 | 55,122 | 59 | 6.38 | 0.01 | 99.89 |
| | Longfin Yellowtail | YTL | *Seriola rivoliana* | LC | 82,108 | 82,087 | 21 | 9.51 | 0.00 | 99.97 |
| Rays | Pelagic stingray | PLS | *Pteroplatytrygon violacea* | LC | 489 | 231 | 258 | 0.03 | 0.04 | 47.24 |

| | | | | | | | | | | |
|---|---|---|---|---|---|---|---|---|---|---|
| | Devil rays | RMM | *Mobula mobular* | EN | 434 | 238 | 196 | 0.03 | 0.03 | 54.84 |
| | | RMT | *Mobula tarapacana* | EN | 195 | 56 | 139 | 0.01 | 0.02 | 28.72 |
| Sharks | Blue shark | BSH | *Prionace glauca* | NT | 456 | 91 | 365 | 0.01 | 0.05 | 19.96 |
| | Silky shark | FAL | *Carcharhinus falciformis* | VU | 23,252 | 16,111 | 7,141 | 1.87 | 0.97 | 69.29 |
| | Scalloped hammerhead | SPL | *Sphyrna lewini* | CR | 1,204 | 1,073 | 131 | 0.12 | 0.02 | 89.12 |
| | Smooth hammerhead | SPZ | *Sphyrna zygaena* | VU | 1,265 | 1,236 | 29 | 0.14 | 0.00 | 97.71 |
| | Shortfin Mako | SMA | *Isurus oxyrinchus* | EN | 131 | 81 | 50 | 0.01 | 0.01 | 61.83 |
| Turtles | Olive ridley turtle | LKV | *Lepidochelys olivacea* | VU | 1,274 | 962 | 312 | 0.11 | 0.04 | 75.51 |
| | Loggerhead turtle | TTL | *Caretta caretta* | VU | 275 | 207 | 68 | 0.02 | 0.01 | 75.27 |

Table S2: Number of bycatch individuals per set, categorized by species group, between 2013 and 2022 on dFADs

| Year | n | Number of sets | Number of other bony fishes per set | Number of billfishes per set | Number of rays per set | Number of sharks per set | Number of turtles per set |
|---|---|---|---|---|---|---|---|
| 2013 | 159,673 | 286 | 556.56 | 0.22 | 0.03 | 1.64 | 0.13 |
| 2014 | 272,389 | 824 | 330.65 | 0.31 | 0.04 | 2.03 | 0.16 |
| 2015 | 604,481 | 896 | 672.18 | 0.29 | 0.02 | 2.12 | 0.05 |
| 2016 | 580,854 | 869 | 664.77 | 0.41 | 0.13 | 2.89 | 0.27 |
| 2017 | 440,571 | 742 | 591.13 | 0.26 | 0.05 | 1.80 | 0.11 |
| 2018 | 633,850 | 1,163 | 542.63 | 0.36 | 0.06 | 1.6 | 0.07 |
| 2019 | 630,614 | 1,089 | 576.36 | 0.3 | 0.08 | 1.9 | 0.12 |
| 2020 | 341,365 | 673 | 505.02 | 0.23 | 0.10 | 1.83 | 0.11 |
| 2021 | 383,301 | 1,011 | 375.74 | 0.27 | 0.03 | 2.99 | 0.14 |
| 2022 | 341,243 | 1,082 | 312.51 | 0.3 | 0.05 | 2.37 | 0.20 |

Table S3: Summary statistics of species-specific catch, including the FAO species code, minimum, maximum, mean, and median number of individuals per set, as well as the minimum and maximum annual mean number of individuals per set. The colours are those of the IUCN protection levels used in table 1.

| FAO | Min ind per set | Max ind per set | Mean ind per set | Median ind per set | Min ind per year | Max ind per year |
|---|---|---|---|---|---|---|
| BUM | 0 | 46 | 0.28 | 0 | 0.17 | 0.39 |
| SAI | 0 | 7 | 0.03 | 0 | 0.01 | 0.06 |
| ALN | 0 | 50 | 0.05 | 0 | 0.01 | 0.17 |
| ALM | 0 | 306 | 0.26 | 0 | 0.09 | 0.84 |
| CNT | 0 | 7,500 | 142.44 | 45 | 104.9 | 229.24 |
| CFW | 0 | 38 | 0.01 | 0 | 0 | 0.08 |
| DIY | 0 | 40 | 0.01 | 0 | 0 | 0.1 |
| DOL | 0 | 752 | 6.34 | 1 | 3.63 | 14.59 |
| EHN | 0 | 23 | 0.01 | 0 | 0 | 0.03 |
| GBA | 0 | 431 | 1.92 | 0 | 1.03 | 3.89 |
| JKY | 0 | 14 | 0.02 | 0 | 0 | 0.04 |
| KYP | 0 | 96 | 0.03 | 0 | 0 | 0.22 |
| KYS | 0 | 455 | 1.51 | 0 | 0.6 | 2.67 |
| LOB | 0 | 369 | 2.15 | 0 | 1.28 | 3.2 |
| MOX | 0 | 3 | 0.01 | 0 | 0.01 | 0.03 |
| MRW | 0 | 4 | 0.01 | 0 | 0 | 0.03 |
| REO | 0 | 99 | 0.03 | 0 | 0 | 0.14 |
| RRU | 0 | 4425 | 91.6 | 20 | 38.7 | 143.56 |
| RUB | 0 | 16,000 | 240.66 | 50 | 104.31 | 361.23 |
| RZV | 0 | 100 | 0.01 | 0 | 0 | 0.37 |
| TRG | 0 | 300 | 1 | 0 | 0.33 | 2.72 |
| USE | 0 | 1,300 | 1.52 | 0 | 0 | 36.79 |
| WAH | 0 | 1,516 | 6.38 | 0 | 5.03 | 8.88 |
| YTL | 0 | 2,250 | 9.51 | 0 | 0.55 | 36.78 |
| PLS | 0 | 11 | 0.03 | 0 | 0.01 | 0.08 |
| RMM | 0 | 14 | 0.03 | 0 | 0 | 0.06 |
| RMT | 0 | 10 | 0.01 | 0 | 0 | 0.03 |
| BSH | 0 | 13 | 0.01 | 0 | 0 | 0.04 |
| FAL | 0 | 185 | 1.87 | 0 | 1.41 | 2.78 |
| SPL | 0 | 33 | 0.12 | 0 | 0 | 0.34 |
| SPZ | 0 | 45 | 0.14 | 0 | 0.02 | 0.39 |
| SMA | 0 | 7 | 0.01 | 0 | 0 | 0.02 |
| LKV | 0 | 7 | 0.11 | 0 | 0.04 | 0.25 |
| TTL | 0 | 5 | 0.02 | 0 | 0 | 0.07 |

Table S4: Number of bycatch caught by species group between 2013 and 2022 on fishing aggregating devices

| Year | n | Number of other bony fishes | Number of billfishes | Number of rays | Number of sharks | Number of turtles |
|---|---|---|---|---|---|---|
| 2013 | 159673 | 159177 | 64 | 8 | 470 | 37 |
| 2014 | 272389 | 270358 | 257 | 37 | 1673 | 130 |
| 2015 | 604481 | 602272 | 259 | 21 | 1897 | 48 |
| 2016 | 580854 | 577687 | 355 | 113 | 2515 | 233 |
| 2017 | 440571 | 438615 | 196 | 36 | 1339 | 83 |
| 2018 | 633850 | 631077 | 419 | 71 | 1850 | 81 |
| 2019 | 630614 | 627659 | 331 | 84 | 2067 | 126 |
| 2020 | 341365 | 339876 | 158 | 66 | 1230 | 73 |
| 2021 | 383301 | 379869 | 269 | 35 | 3021 | 137 |
| 2022 | 341243 | 338140 | 323 | 54 | 2567 | 221 |

Table S5: Size maturity (L50) in centimetres of each threatened species,

| Species group | FAO | | Scientific name | Fishbase (size at Maturity L50) |
|---|---|---|---|---|
| Billfishes | BUM | VU | *Makaira nigricans* | / |
| Other bony fishes | MOX | VU | *Mola mola* | / |
| | TRG | VU | *Balistes capriscus* | / |
| Rays | RMM | EN | *Mobula mobular* | 200 cm WD (McEachran and Séret, 1990) |
| | RMT | EN | *Mobula tarapacana* | 200 cm WD (White et al., 2006) |
| Sharks | BSH | NT | *Prionace glauca* | 221 cm TL (Cervigón et al., 1992; Muus and Nielsen, 1999) |
| | FAL | VU | *Carcharhinus falciformis* | 225 cm TL (Compagno and Niem, 1998) |
| | SMA | EN | *Isurus oxyrinchus* | 280 cm TL (Cervigón et al., 1992; Weigmann, 2016) |
| | SPL | CR | *Sphyrna lewini* | 230 cm TL (Smith, 1997; Compagno, 1998) |
| | SPZ | VU | *Sphyrna zygaena* | 265 cm TL (Compagno, 1998; Muus and Nielsen, 1999) |
| Turtles | LKV | VU | *Lepidochelys olivacea* | 66 cm CCL (Marquez, 1990) |
| | TTL | VU | *Caretta caretta* | 80 cm CCL (Casale et al., 2011) |

Table S6: Catch rates of threatened species in the Atlantic (number of individuals per set) by region

| | **Atlantic** | **Mauritania** | **Gabon** | **Angola** | **Guinea** |
|---|---|---|---|---|---|
| **Number of sets** | 8635 | 248 | 973 | 702 | 2555 |
| Silky shark | **1.87** | 0.33 | **2.81** | **4.21** | **1.28** |
| Billfish | **0.28** | 0.28 | 0.30 | 0.23 | **0.24** |
| Smooth hammerhead | 0.14 | **2.45** | 0.38 | 0.09 | 0.01 |
| Scalloped hammerhead | 0.12 | **0.40** | **0.64** | 0.22 | 0.02 |
| Olive ridley turtle | 0.11 | 0.08 | 0.10 | **0.87** | 0.01 |
| Devil ray | 0.03 | 0.27 | 0.00 | 0.14 | 0.01 |
| Loggerhead turtle | 0.02 | **0.43** | 0.01 | 0.06 | 0.00 |
| Shortfin mako | 0.01 | 0.10 | 0.01 | 0.02 | 0.00 |
| Blue shark | 0.01 | 0.02 | 0.05 | 0.01 | 0.00 |

Table S7: Percentages of adult and juvenile individuals, for each hotspot, broken down into three categories: males, females and total. The hotspots studied are Mauritania, Gabon, Angola and Guinea, as well as an 'Other' category covering the rest of the Atlantic (outside these four hotspots). The results of the Chi² test are shown in coloured boxes: in green when the residuals are greater than 2, indicating significant over-representation compared with the other hotspots; in orange when the residuals are less than -2, indicating significant under-representation.

| | | | % Adult | | | % Juvenile | | |
|---|---|---|---|---|---|---|---|---|
| FAO | Hotspot | Nb. Indiv. Per set | Total | Male | Female | Total | Male | Female |
| FAL | Mauritania | 0.33 | 0 | 0 | 0 | 96 | 56 | 40 |
| | Gabon | 2.81 | 3 | 1 | 2 | 91 | 50 | 41 |
| | Angola | 4.21 | 1 | 0 | 1 | 89 | 41 | 48 |
| | Guinea | 1.28 | 0 | 0 | 0 | 93 | 43 | 50 |
| | Other | 1.87 | 0 | 0 | 0 | 95 | 44 | 52 |
| SPL | Mauritania | 0.40 | 13 | 1 | 12 | 78 | 9 | 69 |
| | Gabon | 0.64 | 29 | 12 | 17 | 65 | 34 | 31 |
| | Angola | 0.22 | 34 | 10 | 24 | 57 | 26 | 31 |
| | Guinea | 0.02 | 12 | 5 | 7 | 84 | 32 | 52 |
| | Other | 0.12 | 46 | 14 | 32 | 54 | 22 | 32 |
| SPZ | Mauritania | 2.45 | 1 | 0 | 1 | 87 | 27 | 60 |
| | Gabon | 0.38 | 3 | 0 | 3 | 90 | 41 | 49 |
| | Angola | 0.09 | 0 | 0 | 0 | 95 | 19 | 76 |
| | Guinea | 0.01 | 22 | 11 | 11 | 78 | 56 | 22 |
| | Other | 0.14 | 6 | 3 | 3 | 91 | 48 | 43 |
| RMM-T | Mauritania | 0.27 | 85 | 36 | 49 | 15 | 4 | 11 |
| | Gabon | 0.00 | 0 | 0 | 0 | 100 | 100 | 0 |
| | Angola | 0.14 | 61 | 44 | 17 | 39 | 12 | 27 |
| | Guinea | 0.01 | 86 | 29 | 57 | 15 | 5 | 10 |
| | Other | 0.03 | 63 | 45 | 18 | 36 | 16 | 20 |
| BSH | Mauritania | 0.02 | 0 | 0 | 0 | 100 | 75 | 25 |
| | Gabon | 0.05 | 60 | 53 | 7 | 40 | 38 | 2 |
| | Angola | 0.01 | 12 | 12 | 0 | 87 | 75 | 12 |
| | Guinea | 0.00 | 22 | 11 | 11 | 67 | 0 | 67 |
| | Other | 0.01 | 69 | 38 | 31 | 31 | 8 | 23 |
| LKV | Mauritania | 0.01 | 91 | 73 | 18 | 9 | 9 | 0 |
| | Gabon | 0.02 | 91 | 53 | 38 | 9 | 3 | 6 |

| | | | | | | | | |
|---|---|---|---|---|---|---|---|---|
| | Angola | 0.14 | 83 | 55 | 28 | 12 | 6 | 6 |
| | Guinea | 0.00 | 65 | 35 | 30 | 34 | 17 | 17 |
| | Other | 0.03 | 73 | 47 | 26 | 25 | 10 | 15 |
| TTL | Mauritania | 0.11 | 5 | 3 | 2 | 93 | 28 | 65 |
| | Gabon | 0.00 | 50 | 10 | 40 | 50 | 20 | 30 |
| | Angola | 0.00 | 6 | 6 | 0 | 83 | 50 | 33 |
| | Guinea | 0.00 | 14 | 14 | 0 | 86 | 29 | 57 |
| | Other | 0.00 | 16 | 10 | 6 | 84 | 45 | 39 |

Table S8: Contingency tables used to compare sex and life-stage composition among bycatch hotspots

| | Mauritania | Gabon | Angola | Guinea | Other |
|---|---|---|---|---|---|
| SPZ_nb_male_juvenile | 130 | 122 | 4 | 5 | 64 |
| SPZ_nb_female_juvenile | 283 | 147 | 16 | 2 | 63 |
| SPZ_nb_male_adult | 0 | 1 | 0 | 1 | 4 |
| SPZ_nb_female_adult | 5 | 8 | 0 | 1 | 6 |
| SPL_nb_male_juvenile | 6 | 159 | 36 | 14 | 19 |
| SPL_nb_female_juvenile | 46 | 147 | 38 | 23 | 33 |
| SPL_nb_male_adult | 1 | 55 | 13 | 2 | 12 |
| SPL_nb_female_adult | 8 | 79 | 33 | 3 | 37 |
| BSH_nb_male_juvenile | 3 | 21 | 6 | 0 | 5 |
| BSH_nb_female_juvenile | 1 | 1 | 1 | 6 | 15 |
| BSH_nb_male_adult | 0 | 29 | 1 | 1 | 13 |
| BSH_nb_female_adult | 0 | 4 | 0 | 1 | 7 |
| RMM_nb_male_juvenile | 2 | 1 | 6 | 1 | 14 |
| RMM_nb_female_juvenile | 5 | 0 | 13 | 2 | 11 |
| RMM_nb_male_adult | 17 | 0 | 21 | 6 | 57 |
| RMM_nb_female_adult | 23 | 0 | 8 | 12 | 47 |
| FAL_nb_male_juvenile | 44 | 929 | 843 | 1047 | 2994 |
| FAL_nb_female_juvenile | 31 | 761 | 1000 | 1200 | 3602 |
| FAL_nb_male_adult | 0 | 26 | 7 | 4 | 38 |
| FAL_nb_female_adult | 0 | 28 | 13 | 5 | 47 |

Table S9: Number of species per set observed in the biodiversity hotspot of Mauritania. Only the months of July to September have enough sets to draw conclusions as they have a total number of sets > 30.

| | **June** | **July*** | **August*** | **September*** | **October** |
|---|---|---|---|---|---|
| **Number of sets** | 11 | 107 | 72 | 49 | 9 |
| Billfishes (BUM) | 0.36 | 0.16 | 0.25 | 0.35 | 1.44 |
| Sunfishes (MOX) | | 0.04 | 0.06 | | |
| Grey Triggerfish (TRG) | | | | 0.02 | |
| Devil Ray (RMM, RMT) | 1.45 | 0.22 | 0.15 | 0.31 | |
| Blue shark (BSH) | | 0.02 | 0.03 | 0.02 | |
| Silky Shark (FAL) | | 0.28 | 0.01 | 0.39 | 3.67 |
| Scalloped hammerhead (SPL) | 0.64 | **0.74** | 0.15 | 0.04 | |
| **Smooth hammerhead (SPZ)** | | **2.29** | **4.69** | **0.49** | |
| Shortfin Mako (SMA) | | 0.13 | 0.12 | 0.04 | |
| Olive ridley turtle (LKV) | 0.09 | 0.08 | 0.03 | 0.18 | |
| Loggerhead turtle (TTL) | 0.27 | **0.42** | 0.31 | **0.71** | 0.11 |

Table S10: Number of species per set observed in the biodiversity hotspot of Gabon. Only the months from May to October have enough sets to draw conclusions as they have a total number of sets > 30.

| | **March** | **May*** | **June*** | **July*** | **August*** | **September*** | **October*** | **November** | **December** |
|---|---|---|---|---|---|---|---|---|---|
| **Number of sets** | 1 | 131 | 193 | 113 | 113 | 191 | 197 | 22 | 10 |
| Billfishes (BUM) | | 0.24 | 0.16 | 0.18 | 0.34 | 0.46 | 0.43 | 0.18 | |
| Sunfishes (MOX) | | 0.02 | 0.01 | 0.02 | 0.02 | 0.04 | 0.01 | | |
| Grey Triggerfish (TRG) | | 0.08 | 0.06 | 0.34 | 0.30 | 0.43 | 0.33 | 0.14 | |
| Devil Ray (RMM, RMT) | | | | | | 0.01 | | | |
| Blue shark (BSH) | | 0.03 | 0.09 | 0.05 | 0.02 | 0.08 | | | |
| **Silky Shark (FAL)** | 1 | **2.58** | **1.59** | **4.05** | **3.08** | **3.47** | **1.84** | **4.45** | 4.40 |
| **Scalloped hammerhead (SPL)** | | 0.52 | 0.21 | 0.55 | **1.49** | **0.79** | 0.08 | | |
| **Smooth hammerhead (SPZ)** | | 0.09 | 0.26 | 0.30 | **1.07** | 0.21 | 0.21 | | |
| Shortfin Mako (SMA) | | 0.02 | 0.01 | 0.02 | | 0.02 | 0.01 | | |
| Olive ridley turtle (LKV) | | 0.02 | 0.04 | 0.15 | 0.21 | 0.12 | 0.02 | 0.09 | 0.10 |
| Loggerhead turtle (TTL) | | | | 0.01 | 0.05 | 0.02 | | | |

Table S11: Number of species per set observed in the biodiversity hotspot of Angola. Only the months from September to November have enough sets to draw conclusions as they have a total number of sets > 30.

| | January | March | May | August | September* | October* | November* | December |
|---|---|---|---|---|---|---|---|---|
| **Number of sets** | 21 | 8 | 3 | 7 | 156 | 430 | 74 | 3 |
| Billfishes (BUM) | 0.29 | | | 0.57 | 0.26 | 0.21 | 0.22 | 1 |
| Sunfishes (MOX) | 0.05 | | | | 0.11 | 0.05 | 0.03 | |
| **Grey Triggerfish (TRG)** | 0.10 | | | | 0.58 | **0.95** | 0.19 | |
| Devil Ray (RMM, RMT) | | | | | 0.03 | 0.11 | 0.04 | |
| Blue shark (BSH) | | 0.12 | | | 0.04 | | | |
| **Silky Shark (FAL)** | 5 | 1.62 | 4.67 | 3.57 | **5.15** | **3.99** | **3.38** | 8.67 |
| Scalloped hammerhead (SPL) | 0.57 | | | | 0.19 | 0.22 | 0.26 | |
| Smooth hammerhead (SPZ) | | 1.88 | | 0.14 | 0.03 | 0.10 | | 0.33 |
| Shortfin Mako (SMA) | | | | 0.14 | 0.05 | 0.02 | | |
| **Olive ridley turtle (LKV)** | 0.10 | 0.25 | | 0.43 | 0.66 | **0.94** | **1.30** | 0.67 |
| Loggerhead turtle (TTL) | 0.10 | | | | 0.13 | 0.04 | 0.01 | |

Table S12: Statistical comparison of sex and life-stage composition of threatened species among bycatch hotspots identified in the Eastern Atlantic, including test type, significance levels, effect sizes (Cramér's V), confidence intervals, and biological interpretation of the magnitude of spatial differences

| Species | Test used | p-value | Cramér's V | 95% CI | Effect size | Interpretation |
|---|---|---|---|---|---|---|
| Rays | Fisher | 0.001 | 0.26 | NA | Moderate | Moderate differences in size–sex composition between hotspots |
| Blue shark | Fisher | <0.001 | 0.45 | NA | Strong | Strong spatial structuring between hotspots |
| Scalloped hammerhead | Chi² | <0.001 | 0.17 | [0.145 – 0.223] | Low–moderate | Weak to moderate differences between hotspots |
| Smooth hammerhead | Fisher | <0.001 | 0.16 | NA | Low | Weak differences; interpretation cautious due to low expected counts |
| Silky shark | Fisher | <0.001 | 0.07 | [0.056 – 0.083] | Very low | Very weak differences; biologically limited |
| Olive ridley turtle | Fisher | 0.010 | 0.12 | [0.096 – 0.189] | Low | Weak differences between hotspots |
| Loggerhead turtle | Fisher | 0.004 | 0.28 | NA | Moderate | Marked structural differences between hotspots |

### Supplementary Figures

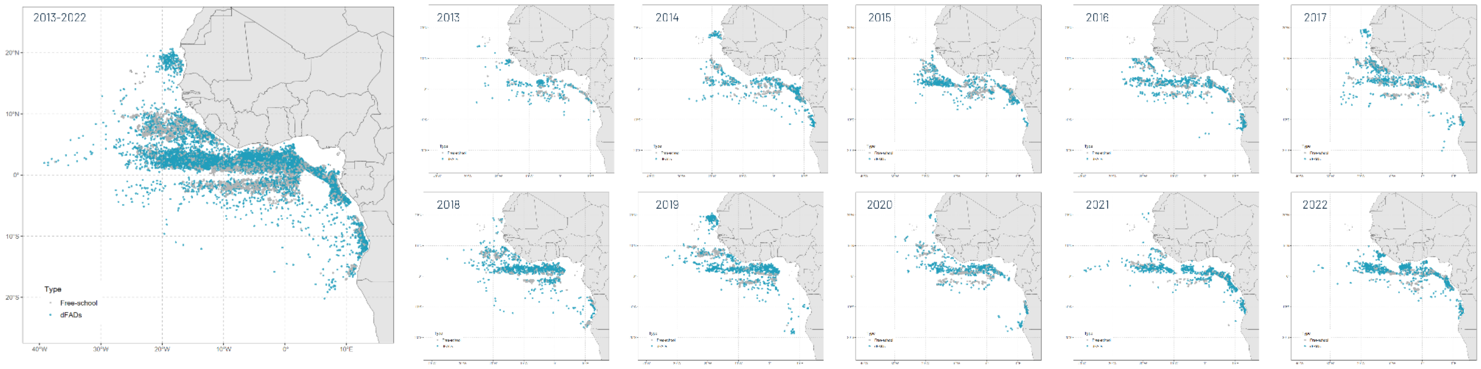


Figure S1. Geographic distribution of purse-seine sets conducted on drifting fish aggregating devices (dFADs; blue point) and free-swimming schools (FSCs; grey point) in the Atlantic Ocean (2013–2022)

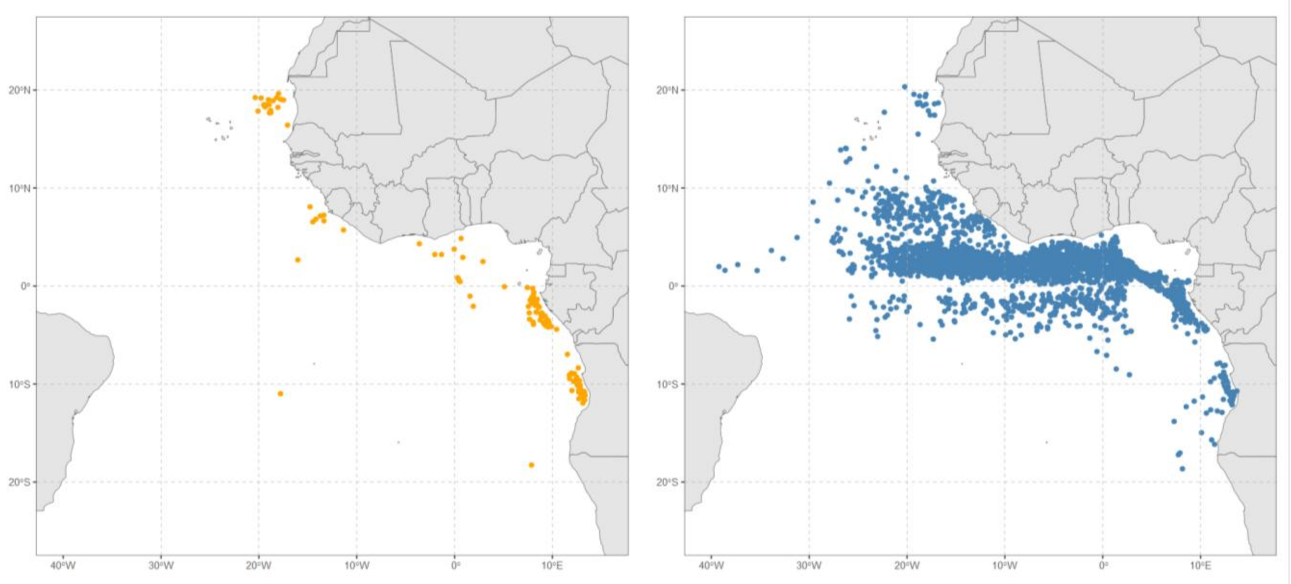


Figure S2:  Geographic occurrence patterns of Cluster A (orange) and Cluster B (blue) species identified from co-occurrence analyses (2013–2022)

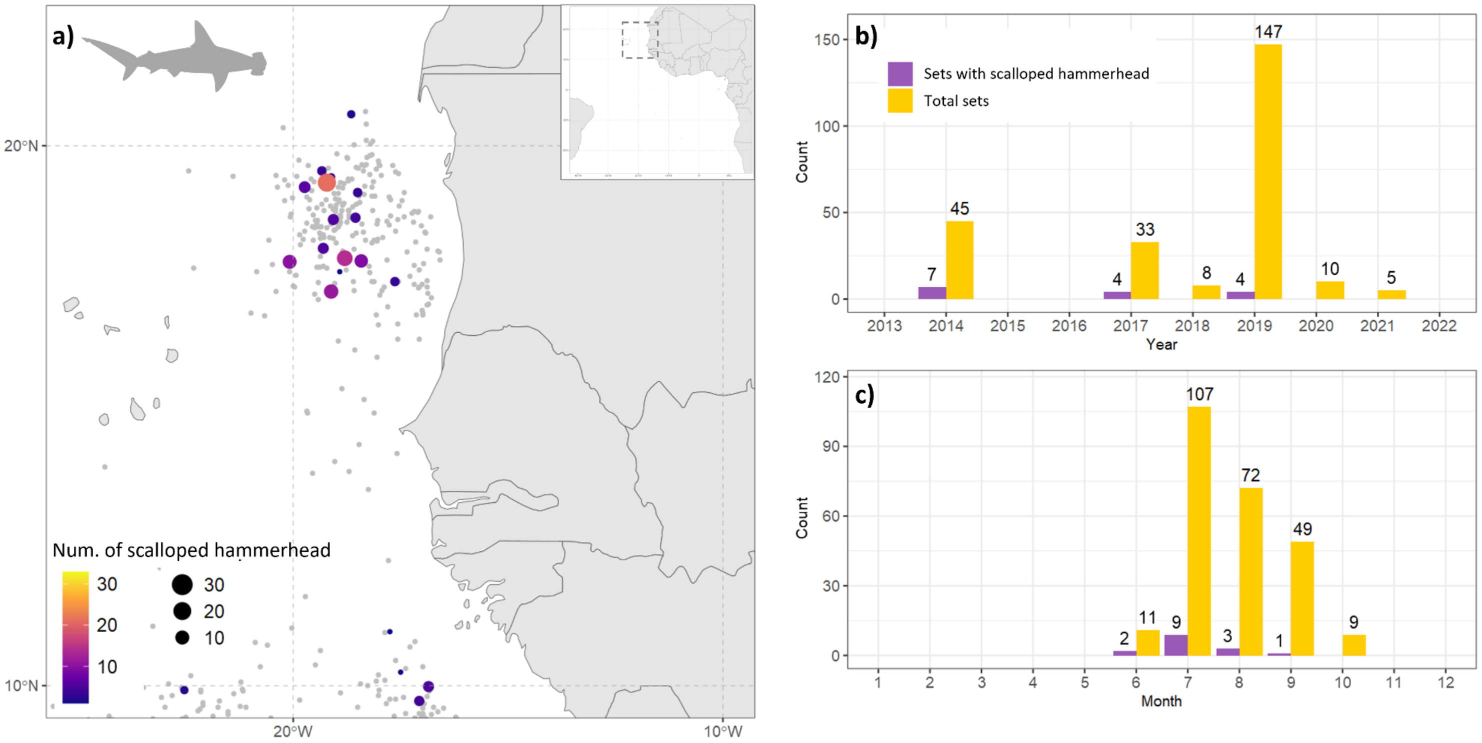


Figure S3: **Spatial and temporal distribution of scalloped hammerhead (SPL).** Map (a) illustrates the locations of scalloped hammerhead captures in Mauritania. The colour and size of the points represent the relative importance of each capture set. The presence of grey points indicates sets that occurred in the region, but did not result in the capture of any scalloped hammerhead individuals. Graphs (b) and (c) illustrate the number of sets with SPL by month and by year, respectively, along with the total number of sets conducted in Mauritania.

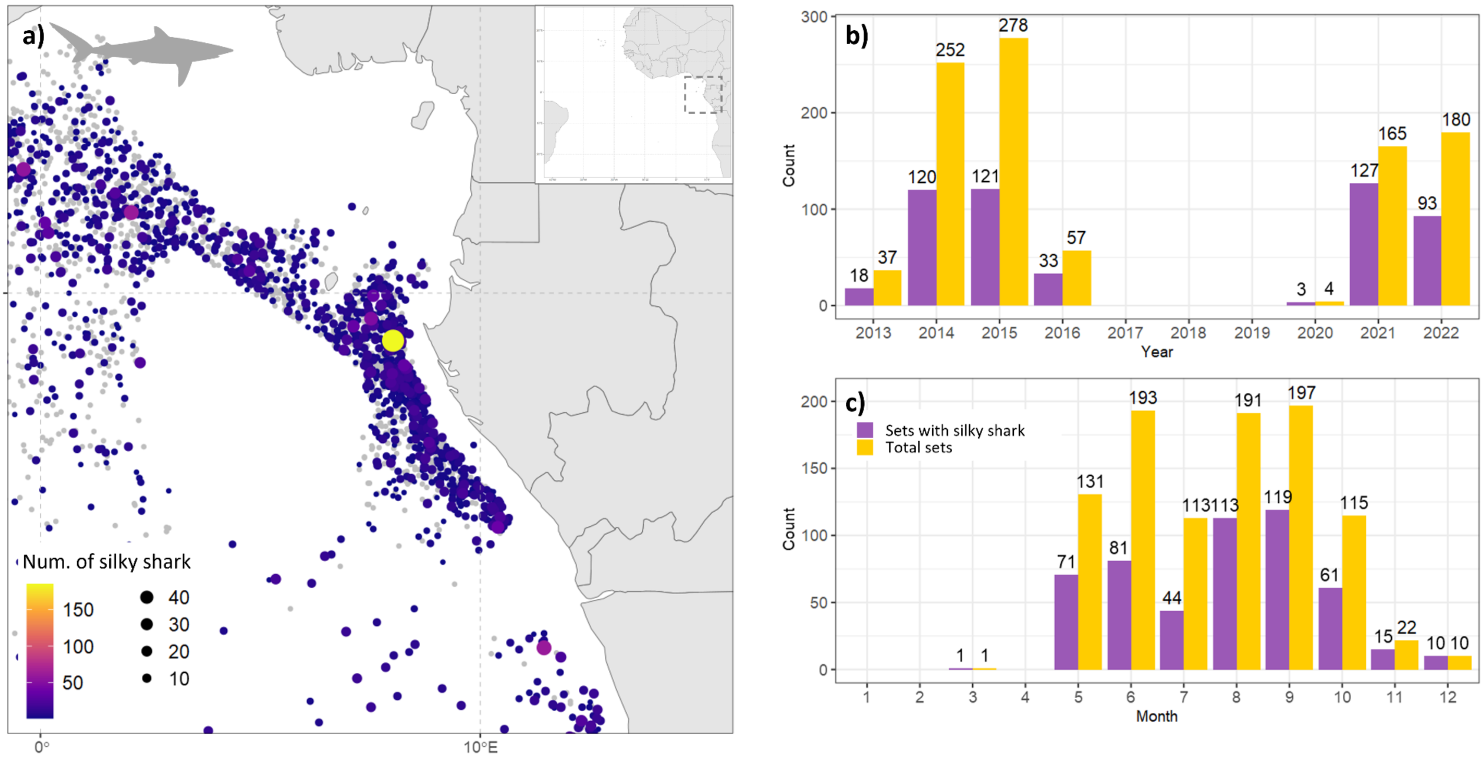


Figure S4: **Spatial and temporal distribution of silky shark (FAL) in Gabon.** Map (a) illustrates the locations of silky shark captures. The colour and size of the points represent the relative importance of each capture set. The presence of grey points indicates sets that occurred in the region, but did not result in the capture of any silky shark individuals. Graphs (b) and (c) illustrate the number of sets with FAL by month and by year, respectively, along with the total number of sets conducted in Gabon.

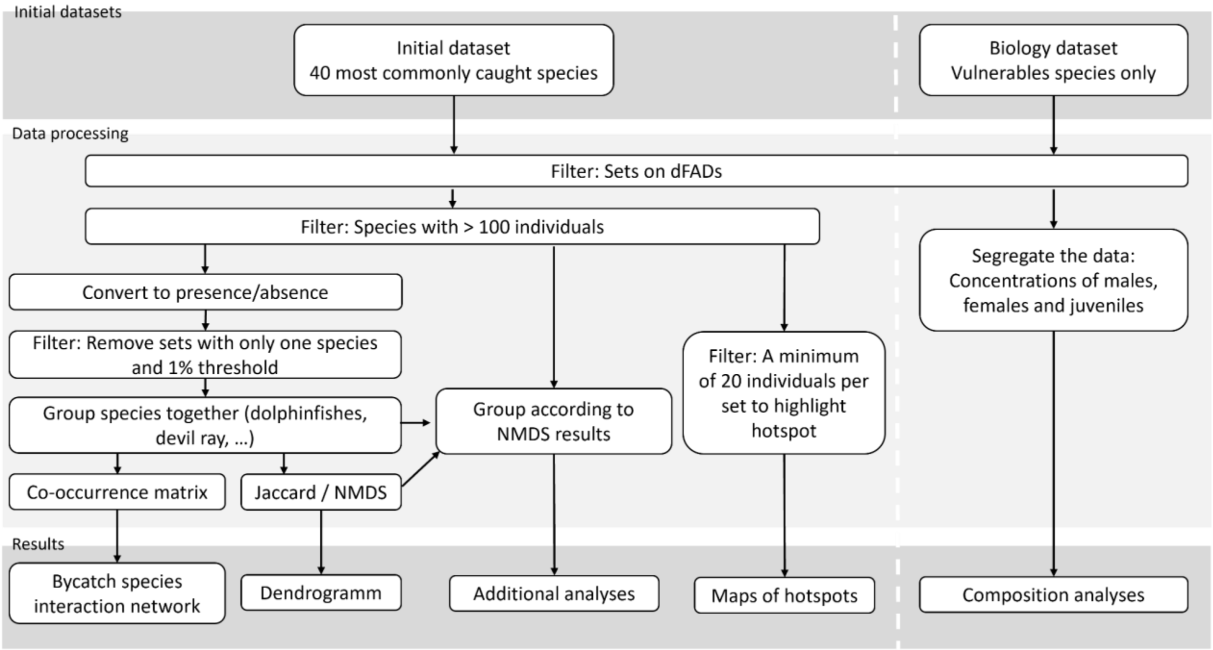


Figure S5: Flowchart of dataset construction and filtering steps used in the analyses.

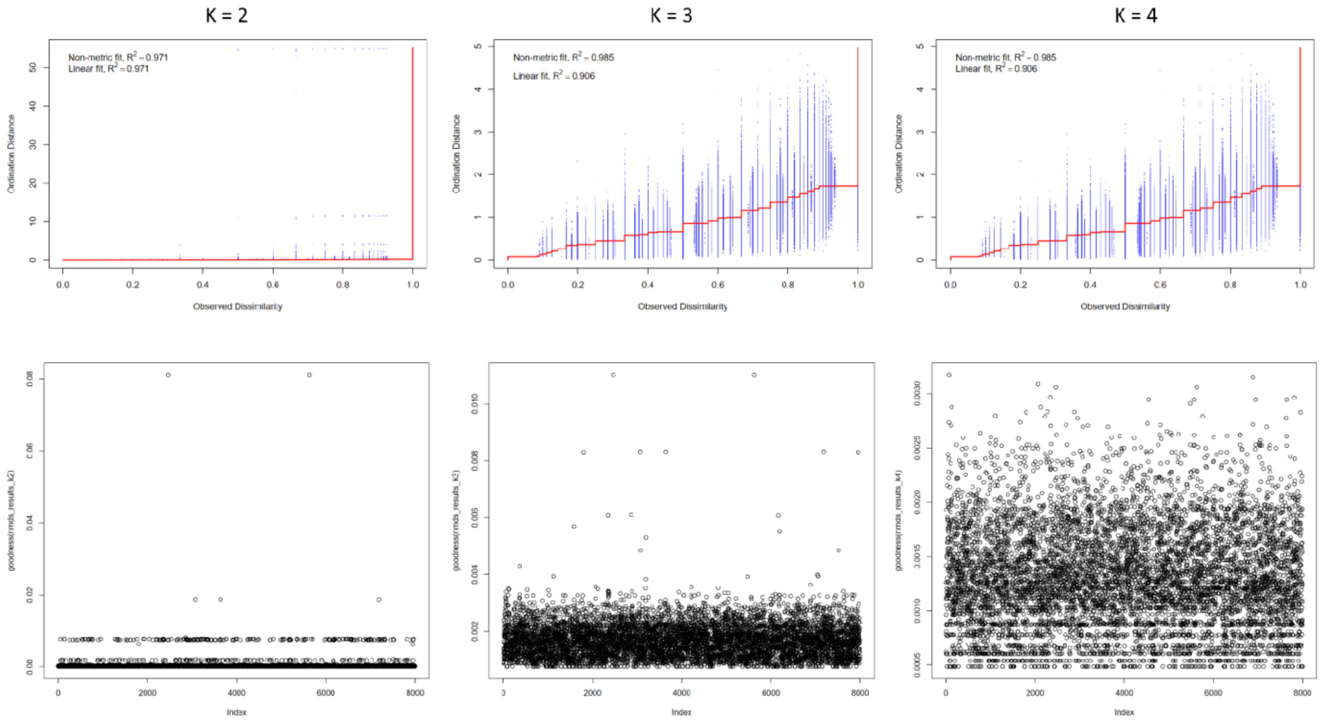


Figure S6: Evaluation of NMDS dimensionality using stress values, Shepard plots, and goodness-of-fit metrics (k = 2–4).

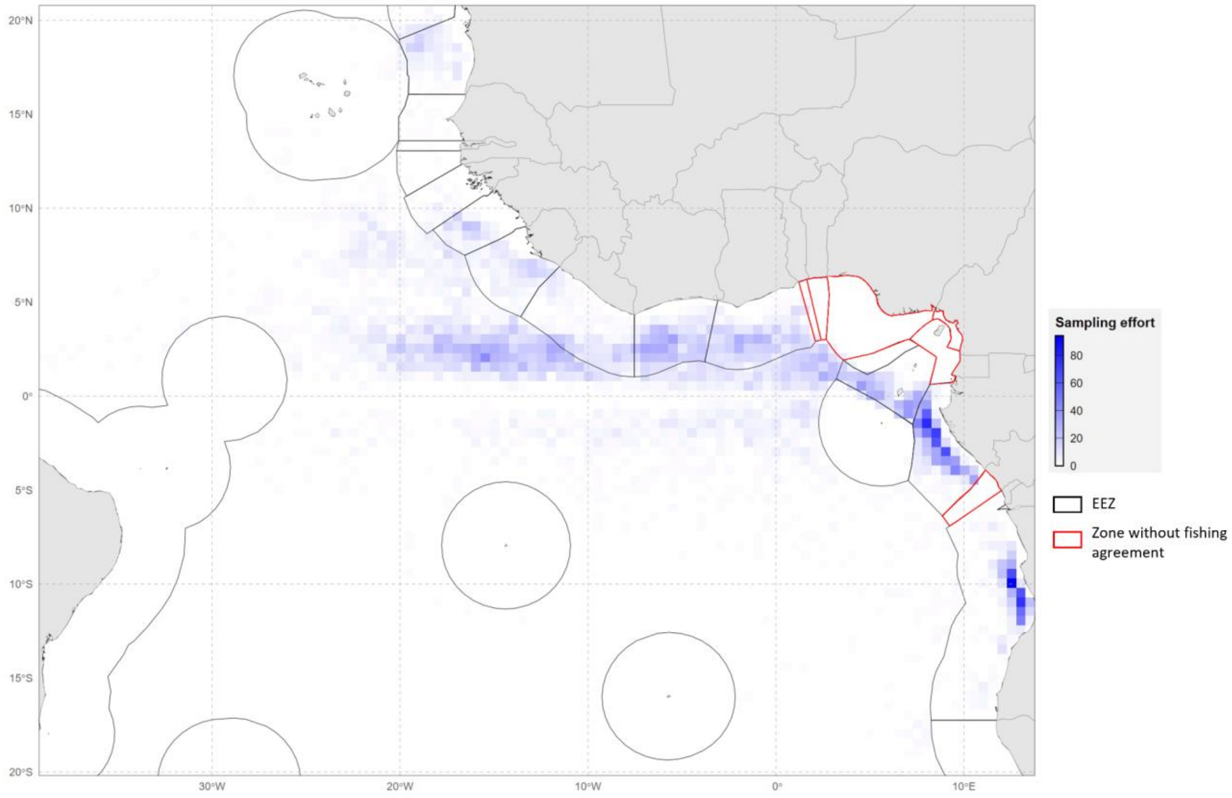


Figure S7: Spatial distribution of purse-seine fishing effort in the Atlantic Ocean, showing Exclusive Economic Zones (EEZs) and areas without fishing agreements (2013–2022). Figure from Lerebourg et al. 2026.
